# Proteins differentially expressed between pathogenic and non-pathogenic *Entamoeba histolytica* clones influence pathogenicity by different mechanisms

**DOI:** 10.1101/2023.06.29.547007

**Authors:** Juliett Anders, Constantin König, Corinna Lender, Arne Hellhund, Sarah Nehls, Ibrahim Shalabi, Barbara Honecker, Stephan Lorenzen, Martin Meyer, Jenny Matthiesen, Dániel Cadar, Thomas Roeder, Nahla Galal Metwally, Hannelore Lotter, Iris Bruchhaus

**Author notes:** Contributed equally (Juliett Anders, Constantin König, Corinna Lender).

## Abstract

Recently, two genes involved in pathogenicity in a mouse model of amoebic liver abscess were identified based on their differential expression between non-pathogenic (A1^np^) and pathogenic (B2^p^) clones of the *Entamoeba histolytica* isolate HM:1-IMSS. While overexpression of a gene encoding the metallopeptidase EhMP8-2 decreases the virulence of the pathogenic clone B2^p^, overexpression of the gene *ehi_127670* (*ehhp127*), encoding a hypothetical protein, increases the virulence of the non-pathogenic clone A1^np^, while silencing this gene in B2^p^ decreases virulence. To understand the role of both molecules in determining the pathogenicity of *E. histolytica*, silencing and overexpression transfectants were characterized in detail. Silencing of *ehmp8-2*, of the homologous gene *ehmp8-1*, or of both together in A1^np^ trophozoites significantly altered the transcript levels of 60-350 genes. This strong change in the expression profile caused by the silencing of *ehmp8-1* and/or *ehmp8-2* implies that these peptidases regulate expression of numerous genes. Consequently, numerous phenotypic characteristics including cytopathic, hemolytic and cysteine peptidase activity were changed in response to their silencing. Silencing of *ehhp127* in B2^p^ trophozoites did not affect other genes, whereas overexpression in A1^np^ trophozoites results in an altered expression of approximately 140 genes. EhHP127 appears to be important for trophozoite movement, as silencing negatively affects and overexpression positively affects trophozoite motility. Interestingly, the specific silencing of *ehhp127* also impairs cytopathic activity, cysteine peptidase and hemolytic activity. All three molecules of interest, namely EhMP8-1, EhMP8-2, and EhHP127 can be detected in amoeba vesicles. Our results clearly show that the proteins studied here influence the pathogenicity of amoebae in different ways and use entirely different mechanisms to do so.

**Author summary:** The human pathogen *Entamoeba histolytica* can live asymptomatically in the intestine or become invasive and cause fatal liver abscesses. Approximately 15,000 people die each year as a result of an amoebic infection. Recently, two clones with different pathogenicity (A1^np^: non-pathogenic; B2^p^: pathogenic) derived from the *E. histolytica* isolate HM:1-IMSS were compared at the transcriptome level. Two highly differentially expressed genes (*ehhp127* encoding a hypothetical protein and *ehmp8-2* encoding a metallopeptidase) were identified. Analysis of *E. histolytica* transfectants showed that silencing of *ehhp127* and overexpression of *ehmp8-2* in B2^p^ trophozoites reduced amoebic liver abscess formation in the mouse model. In this study, we characterized *E. histolytica* silencing and overexpression transfectants of *ehmp8-2*, as well as of the homologous gene *ehmp8-1* and of *ehhp127*. It was shown that the altered expression of the metallopeptidase genes has a strong influence on the expression of a large number of genes and that the phenotype is strongly altered as a result. Silencing of *ehhp127* does not affect the overall expression profile. However, specific silencing has a negative effect on motility, cysteine peptidase, hemolytic and cytopathic activity. All three molecules were shown to be localized in trophozoite vesicles.

## Introduction

*Entamoeba histolytica* is the causative agent of amoebiasis, which kills approximately 15,000 people annually [1]. *E. histolytica* is an intestinal protozoan that, for reasons that are not yet understood, can become invasive, penetrate the intestinal mucosa, invade the tissue, and migrate via the bloodstream to the liver. This invasion can lead to the development of amoebic colitis and the formation of amoebic liver abscesses (ALAs).

In order to identify the virulence factors of *E. histolytica* involved in the formation of ALAs, in recent years we have analyzed amoeba clones (A1^np^ and B2^p^) originally derived from the same isolate (HM- 1:IMSS) but differing in their ability to form ALAs [2-5]. Seven days after injection of amoebae into the liver of mice, no ALAs can be detected in animals infected with A1^np^ trophozoites, whereas ALAs are present in animals infected with B2^p^ trophozoites [4].

Analysis of the transcriptomes of the non-pathogenic clone A1^np^ and the pathogenic clone B2^p^ revealed 76 genes that are differentially expressed between the two clones. These include 46 genes that are significantly more highly expressed in A1^np^ trophozoites and 30 genes that are significantly more highly expressed in B2^p^ trophozoites (fold change >3, padj <0.05) [4].

The second most highly differentially expressed gene in A1^np^ trophozoites is *ehi_042870* (fold change 149), which encodes the cell surface protease gp63 (metallopeptidase EhMP8-2) [4]. The up-regulation of *ehmp8-2* in B2^p^ trophozoites leads to a loss of virulence of these transfectants, thus they are no longer able to form ALAs [4]. Silencing of *ehmp8-2* expression does not alter the non-pathogenic phenotype of A1^np^ [5]. Recently, the genome of *E. histolytica* was analyzed for peptidase-encoding genes. A total of 79 peptidase-encoding genes were identified. The largest group with 45 genes encodes cysteine peptidases (CPs). This is followed by metallopeptidase-encoding genes (21 members), serine peptidase-encoding genes (9 members) and aspartate peptidase-encoding genes (4 members) [6]. The 21-member group of metallopeptidases consists of 11 families, including two members of the M8 (leishmanolysin/gp63 like) family (EhMP8-1 (EHI_200230) and EhMP8-2 (EHI_042870)) [6]. Of all the metallopeptidases, only EhMP8-1 has been characterized to date [7].

EhMP8-1 was detected in vesicles in all trophozoites examined. Furthermore, the protein was also detected on the cell surface in some of the trophozoites [7]. Silencing of *ehmp8-1* resulted in increased cytoadhesion to cell monolayers, decreased cytopathic activity and motility, and increased phagocytosis [7]. EhMP8-1 and EhMP8-2 contain the conserved HEXXH motif of the M8 family and the conserved C-terminal amino acids histidine and methionine [8]. EhMP8-1 consists of 643 amino acids, a putative signal peptide of 15 amino acids, and a putative transmembrane domain at position 605- 627. EhMP8-2 consists of 662 amino acids, a putative signal peptide of 16 amino acids, and a putative transmembrane domain at position 598-620. Both metallopeptidases share only 32 % sequence identity (49 % similarity) (AmoebaDB, release 61, 15 Dec 2022). While *ehmp8-2* is differentially expressed between A1^np^ and B2^p^, as mentioned above, this is not the case for *ehmp8-1* [4]. Interestingly, there is only one member of the M8 family in the human non-pathogenic species *E. dispar*, which is 92 % identical to the *E. histolytica* metallopeptidase EhMP8-2 [6]. In contrast, pathogenic amoebae from reptiles (*E. invadens*) and macaques (*E. nutalli*) contain the two EhMP8 homologs EIN154240/EIN174510 and ENU1_098770/ENU1_206570, respectively (AmoebaDB, 5elease 61, 15 Dec 2022). It can therefore be speculated that lower virulence correlates with the presence or the expression of an *ehmp8-2* homologue.

The most differentially expressed gene (*ehi_127670*) of the pathogenic clone B2^p^ compared to the non-pathogenic clone A1^np^ encodes a hypothetical protein (EhHP127). It is 193-fold more highly expressed in B2^p^ trophozoites than in A1^np^ trophozoites [4]. Increased expression (16.1-fold) was also shown in isolate G3 (*amoebapore*-silenced) compared to the HM-1:IMSS isolate (primary contact: Carol Gilchrist, University of Virginia, School of Medicine; source version: 2011-10-06; release # / date: AmoebaDB rel. 1.0, 2005-JAN-01). In addition, increased expression of *ehhp127* was detected after adaptation of *E. histolytica* to 2 μM auranofin. However, this adaptation leads to the regulation of the expression of several hundred genes, suggesting that it is a very complex adaptation mechanism. Auranofin is an antirheumatic drug that targets the mammalian thioredoxin reductase (TrxR), but is also highly effective against a variety of pathogenic bacteria and protozoan parasites [9]. Recently, a study analyzed the expression profile of *E. histolytica* isolated from the clinical specimens of three patients (asymptomatic carrier: isolate Ax11, patient with amoebic colitis: isolate Ax22; patient with ALA: isolate Ax19). Expression of *ehhp127* was detected in the isolate Ax19 from a patient with ALA, whereas no expression was detected in the isolates Ax11 and Ax22 [10].

Overexpression of *ehhp127* in non-pathogenic A1^np^ trophozoites resulted in ALA formation in 4 out of 9 infected mice, which was nevertheless not significant [4]. On the other hand, silencing of *ehhp127* in the pathogenic B2^p^ trophozoites significantly reduced the sizes of the ALA [5].

In this study, we aim to understand the functions of EhMP8-1, EhMP8-2, and EhHP127 using silencing and overexpression transfectants and to determine their effects on virulence development using various *in vitro* assays. Our results show that silencing of *ehmp8-1* and/or *ehmp8-2* in A1^np^ trophozoites as well as overexpression of *ehhp127* in A1^np^ trophozoites affects the expression of a large number of genes, whereas silencing of *ehhp127* in B2^p^ trophozoites has no effect on the expression profile of other genes. It is therefore not surprising that silencing of metallopeptidase-encoding genes affects several phenotypic characteristics of amoebae, such as cytopathic, cysteine peptidase and hemolytic activities. Motility is impaired in both overexpression and silencing EhHP127 transfectants, while silencing also has a negative effect on cytopathic, cysteine and hemolytic activity. The metallopeptidases and EhHP127 have in common that they can be detected in trophozoite vesicles. In conclusion, the pathogenicity factor EhHP127 and the metalloproteinase EhMP8-2, which dampens pathogenicity employ different mechanisms to induce their phenotypes.

## Results

### Silencing and overexpression of *ehmp8-1* and *ehmp8-2* in pathogenic and non-pathogenic amoebae

Recently, *ehmp8-1* was shown to be expressed in the non-pathogenic clone A1^np^ at almost the same level as in the pathogenic clone B2^p^ (reads: 2028 vs 2228; [4]). The expression level of *ehmp8-2* in clone A1^np^ is only about half that of *ehmp8-1* (reads: 1098), while *ehmp8-2* is almost not expressed at all in clone B2^p^ (reads: 7.4) [4].

To characterize the metallopeptidases, A1^np^ transfectants were generated in which the expression of *ehmp8-1* (A1^np^MP8-1^Si^), *ehmp8-2* (A1^np^MP8-2^Si^) or both (A1^np^MP8-1+2^Si^) was silenced. Furthermore, B2^p^ transfectants were generated in which *ehmp8-1* (B2^p^MP8-1^Si^) was silenced and *ehmp8-2* (B2^p^MP8-2^OE^) was overexpressed. Real-time quantitative PCR (qRT-PCR) confirmed the successful silencing and overexpression of the corresponding genes in the transfectants (Fig 1A and 1B).

**Fig 1.**
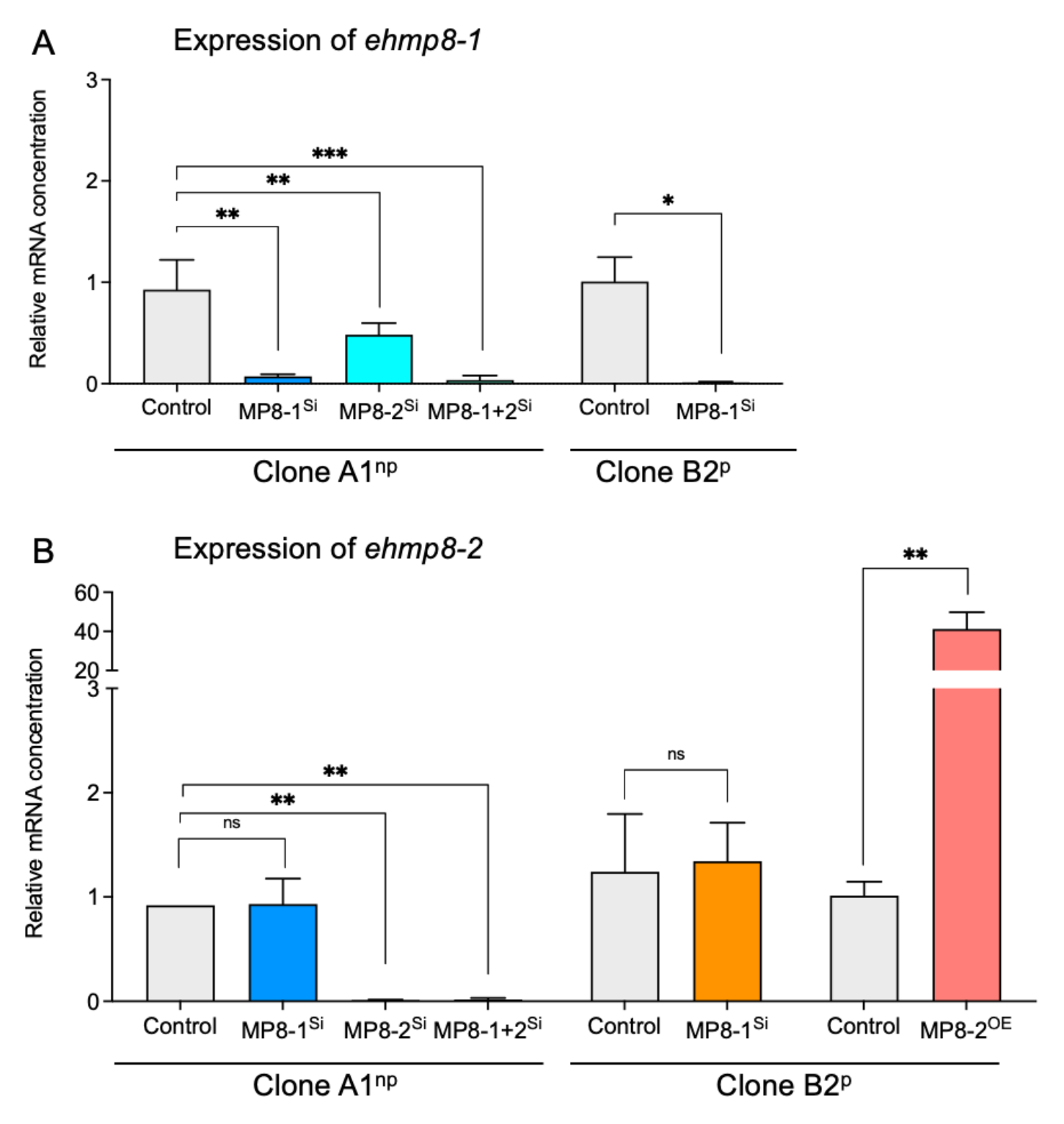
mRNA expression profile of *ehmp8-1* (A) and *ehmp8-2* (B) in A1^np^ and B2^p^ silencing and overexpression transfectants by means of RT-qPCR. RNA isolated from trophozoites was transcribed into cDNA and used for qPCR with SYBR Green to determine the relative mRNA concentration of *ehmp8-1* (A) and *ehmp8-2* (B) of the silencing transfectants A1^np^MP8-1^Si^, A1^np^MP8-2^Si^, A1^np^MP8-1+2^Si^, B^p^MP8-1^Si^ and the overexpression transfectant B2^p^MP8-2^OE^. Actin was used as a calibrator and controls were normalized to 1. n=2-4 (in duplicate). ns: not significant, \**p*<0.05, \*\**p*<0.01, \*\*\**p*<0.001 (Mann-Whitney *U* test).

The transcriptomes of the transfectants were then analyzed. This showed that silencing had a significant effect on the expression of other genes (padj <0.05, fold change >1.8). Silencing of *ehmp8-1* in clone A1^np^ (A1^np^MP8-1^Si^ transfectant) resulted in altered expression of 216 genes (38 upregulated, 178 downregulated), silencing of *ehmp8-2* (A1^np^MP8-2^Si^ transfectant) resulted in altered expression of 347 genes (70 upregulated, 277 downregulated), and silencing of both metallopeptidase genes (A1^np^MP8-1+2^Si^ transfectant) resulted in altered expression of 58 genes (12 upregulated, 46 downregulated) (Fig 2; Fig 3A-3C, S1-S3 Tables). Depending on the transfectant, between 3.8 and 4.7 times more genes were downregulated than upregulated. Here, about 47 % of the genes encode hypothetical proteins, making it difficult to conclude whether specific pathways are affected by the silencing of metallopeptidase genes (S1-S3 Tables).

**Fig 2.**
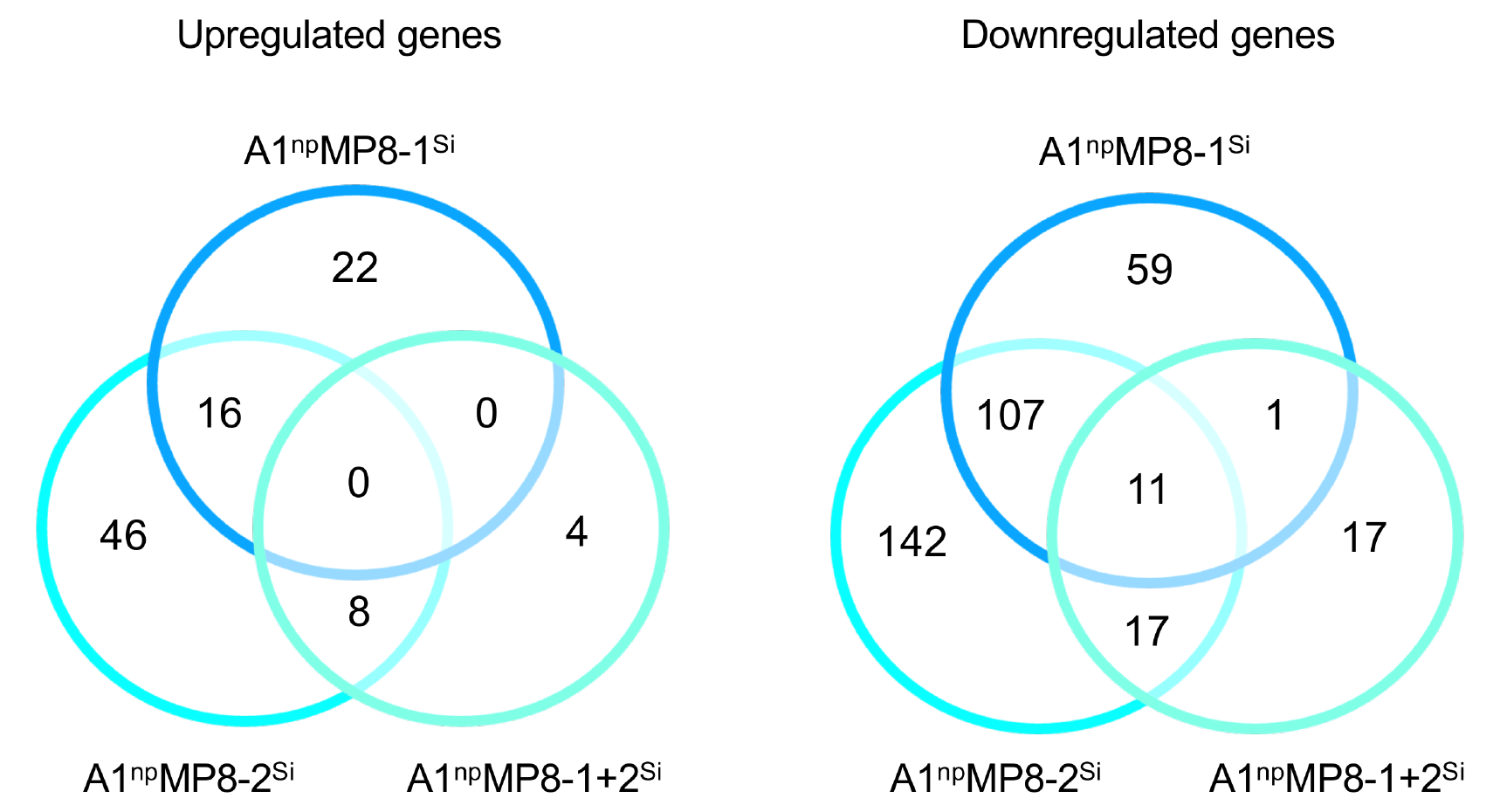
Venn diagram of differentially expressed genes in metallopeptidase silencing transfectants of non-pathogenic clone A1^np^ (A1^np^MP8-1^Si^, A1^np^MP8-2^Si^, A1^np^MP8-1+2^Si^). Padj <0.05, fold change >1.8.

**Fig 3.**
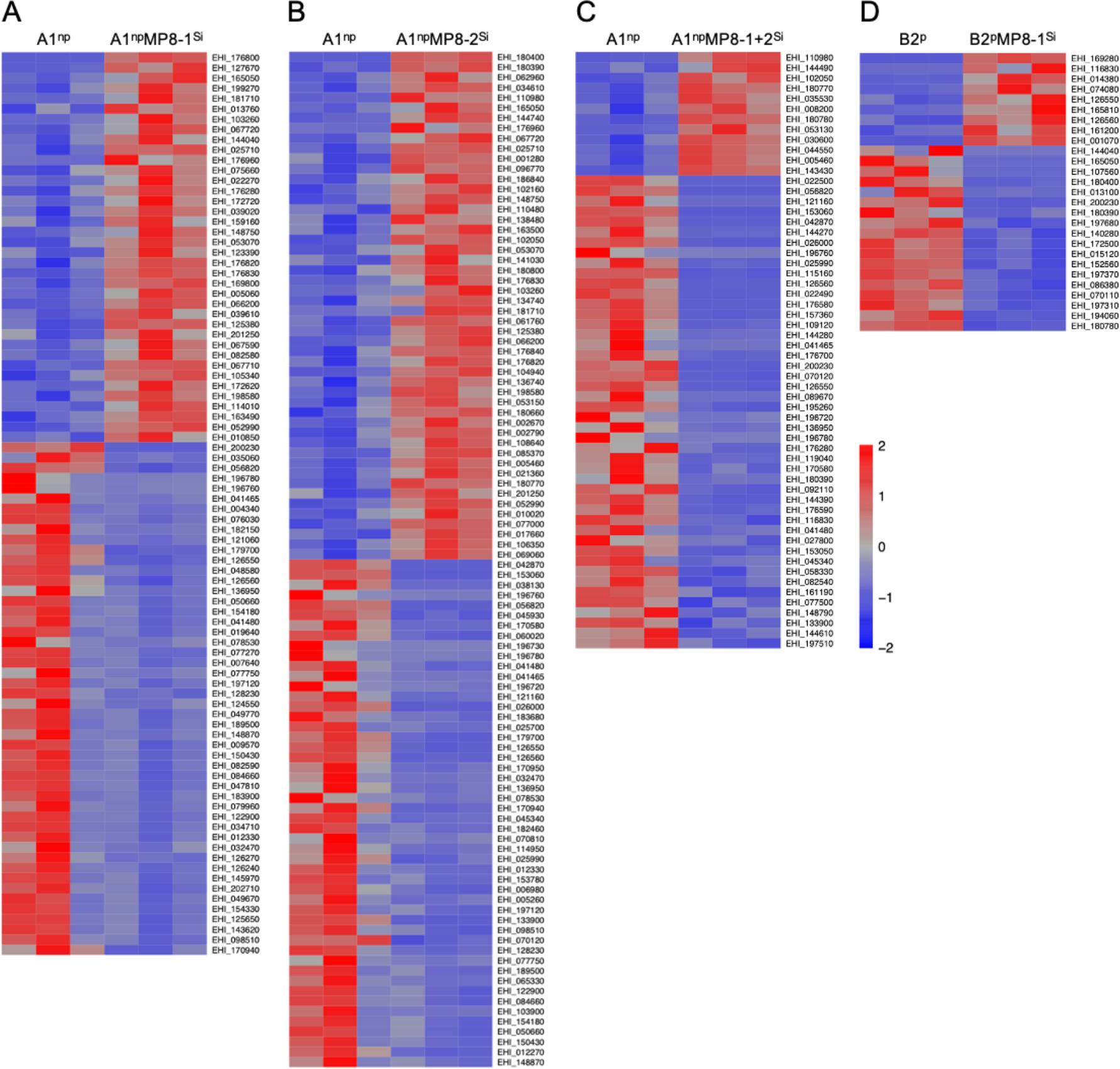
Heatmap of significantly differentially expressed genes (>1.8 fold, padj<0.05) in the different *E. histolytica* transfectants silencing or overexpressing *ehmp8-1*, *ehmp8-2* or both compared to the corresponding controls. A maximum of 50 genes with the highest fold change are shown. A. A1^np^ versus A1^np^MP8-1^Si^, B. A1^np^ versus A1^np^MP8-2^Si^, C. A1^np^ versus A1^np^MP8-1+2^Si^, D. B2^p^ versus B2^p^MP8-1^Si^.

Looking at the genes individually, *ehhp127* was found to be the second most overexpressed gene in A1^np^EhMP8-1^Si^ transfectants (22-fold change, padj 1,67E-19; Fig 3A). Furthermore, the expression of genes encoding proteins that protect against oxidative stress, such as peroxiredoxin, NADPH-dependent FMN reductase domain-containing protein, NADPH-dependent FMN reductase domain-containing protein, and iron-containing superoxide dismutase, are significantly upregulated. In addition, genes encoding the peptidases CAAX prenyl protease and EhCP-A7 are significantly upregulated whereas *ehcp-a4* is downregulated (S1 Table). The 15 most downregulated genes also included 4 *aig* genes (between 5.3- and 6.7-fold) (Table 1).

**Table 1.** *Aig1* genes with significantly differential expression in A1^np^MP8-1^Si^, A1^np^MP8-2^Si^, A1^np^MP8-1+2^Si^, B2^p^MP8-1^Si^, A1^np^HP127^OE^ transfectants compared to respective controls.

| *Aig1* gene<br>(Accession<br>number) | A1<sup>np</sup>MP8-1<sup>Si</sup> |  | A1<sup>np</sup>MP8-2<sup>Si</sup> |  | A1<sup>np</sup>MP8-1+2<sup>Si</sup> |  | B2<sup>p</sup>MP8-1<sup>Si</sup> |  | A1<sup>np</sup>HP127<sup>OE</sup> |  |
| --- | --- | --- | --- | --- | --- | --- | --- | --- | --- | --- |
|  | UP<sup>1</sup> | DOWN<sup>2</sup> | UP | DOWN | UP | DOWN | UP | DOWN | UP | DOWN |
| EHI_176280 | X |  |  |  |  |  |  |  |  | X |
| EHI_136950 |  | X |  | X |  |  |  |  |  |  |
| EHI_126560 |  | X |  | X |  |  | X |  |  | X |
| EHI_126550 |  | X |  | X |  | X | X |  |  | X |
| EHI_041465 |  | X |  |  |  |  |  |  |  |  |
| EHI_180390 |  |  | X |  |  |  |  |  |  |  |
| EHI_026000 |  |  |  | X |  |  |  |  |  |  |
| EHI_025990 |  |  |  | X |  |  |  |  |  |  |
| EHI_176700 |  |  |  | X |  | X |  |  |  | X |
| EHI_022490 |  |  |  | X |  | X |  |  |  |  |
| EHI_041465 |  |  |  | X |  | X |  |  |  |  |
| EHI_089670 |  |  |  |  |  | X |  |  |  | X |
| EHI_157360 |  |  |  |  |  | X |  |  |  | X |
| EHI_144270 |  |  |  |  |  | X |  |  |  |  |
| EHI_176590 |  |  |  |  |  |  |  |  |  | X |
| EHI_176580 |  |  |  |  |  |  |  |  |  | X |
| EHI_109120 |  |  |  |  |  |  |  |  |  | X |
| EHI_144270 |  |  |  |  |  |  |  |  |  | X |

<sup>1</sup>Significant upregulation of expression <sup>2</sup>Significant downregulation of expression (padj<0.05).

The second most upregulated gene in the A1^np^EhMP8-2^Si^ transfectant (EHI_180390; fold change 8.7, padj 4.9E-10) also encodes a protein of the AIG1 family (Table 1, Fig 3B, S2 Table). However, a total of 8 genes encoding AIG1 family proteins are downregulated (Table 1, S2 Table). In addition, as in A1^np^EhMP8-1^Si^ transfectants, a number of genes that could encode for antioxidants (NADPH-dependent FMN reductase domain containing protein, iron-sulfur flavoprotein, iron hydrogenase, peroxiredoxin) are upregulated. It is also striking that many genes encoding proteins related to the cytoskeleton and surface proteins are downregulated (surface antigen ariel1, myotubularin, calponin homology domain protein, formin homology 2 family protein, caldesmon, actin-binding protein, Gal/GalNAc lectin 35 kDa subunit, Gal/GalNAc lectin 170 kDa subunit, filopodin, villin, formin homology 2 family protein, villidin, actinin-like protein) (S2 Table).

Looking at the A1^np^ silencing transfectants in which both metallopeptidase genes were silenced (A1^np^EhMP8-1+2^Si^), it was surprising that silencing had the least effect on the expression and that few of them were found in the intersection of all three transfectants. However, it was very striking that 20 of the 48 downregulated genes belonged to the *aig1* family (Table 1, Fig 3C, S3 Table). In addition, the genes encoding the large and small subunits of the Gal/GalNAc lectin and the cysteine peptidase EhCP-B2 were also downregulated (S3 Table).

Silencing of *ehmp8-1* in the pathogenic clone B2^p^ regulates the expression of only a small number of genes. The expression of a total of 27 genes was significantly affected, of which 9 were upregulated and 18 were downregulated in their expression. Again, two of the 9 upregulated genes are *aig1* genes, while one *aig1* gene was downregulated (Table 1, Fig 3D, S4 Table).

### Phenotypic characteristics of metallopeptidase transfectants

To obtain information on the effect of silencing and overexpression on the phenotype of *E. histolytica,* the motility, phagocytosis rate, cytopathic, cysteine peptidase and hemolytic activities of the transfectants were analyzed. Although silencing affects the expression of a number of genes, it has only little effect on amoeba motility and phagocytosis. Significantly increased motility (*p*=0.0043) was only observed in A1^np^EhMP8-2^Si^ transfectants (Fig 4A) and significantly increased erythrophagocytosis (*p*=0.0212) was observed only in A1^np^EhMP8-1^Si^ transfectants (Fig 4B). The situation was different for the cytopathic activity (monolayer destruction), cysteine peptidase activity and hemolytic activity. Silencing of *ehmp8-1* or *ehmp8-2* alone in both clones A1^np^ and B2^p^ resulted in significant reduction of the cell monolayer destruction between 33 % – 53 % (A1^np^EhMP8-1^Si^, *p*=0.0005; A1^np^EhMP8-2^Si^, *p*=0.0003, B2^p^EhMP8-1^Si^, *p*<0.0001) (Fig 4C and 4H). However, silencing of both metallopeptidase genes had no effect on cytopathic activity (Fig 4C). Surprisingly, overexpression of *ehmp8-2* in B2^p^ trophozoites (B2^p^MP8-2^OE^) also led to significant reduction in monolayer destruction (*p*=0.0012) (Fig 4M). Furthermore, silencing led to a significant reduction in cysteine peptidase activity in all silencing transfectants examined (A1^np^EhMP8-1^Si^, p=0.0052; A1^np^EhMP8-2^Si^, *p*<0.0001; A1^np^EhMP8-1+2^Si^, *p*<0.0001; B2^p^EhMP8-1^Si^, *p*=0.0010). The cysteine peptidase activity in A1^np^ transfectants decreased from 10.3±1.4 (control) to 7.4±2.7 mU/mg (A1^np^EhMP8-1^Si^), 4.3±1.9 mU/mg (A1^np^EhMP8-2^Si^), and 6.2±2.4 mU/mg (A1^np^EhMP8-1+2^Si^) (Fig 4D). The cysteine peptidase activity of B2^p^ trophozoites was 60±15 mU/mg and decreased to 43±11 mU/mg after silencing of *ehmp8-1* (Fig 4I). In contrast, overexpression of *ehmp8-2* in B2^p^ trophozoites resulted in a significant increase in cysteine peptidase activity (B2^p^MP8-2^OE^, p=0.0004) (Fig 4N).

**Fig 4.**
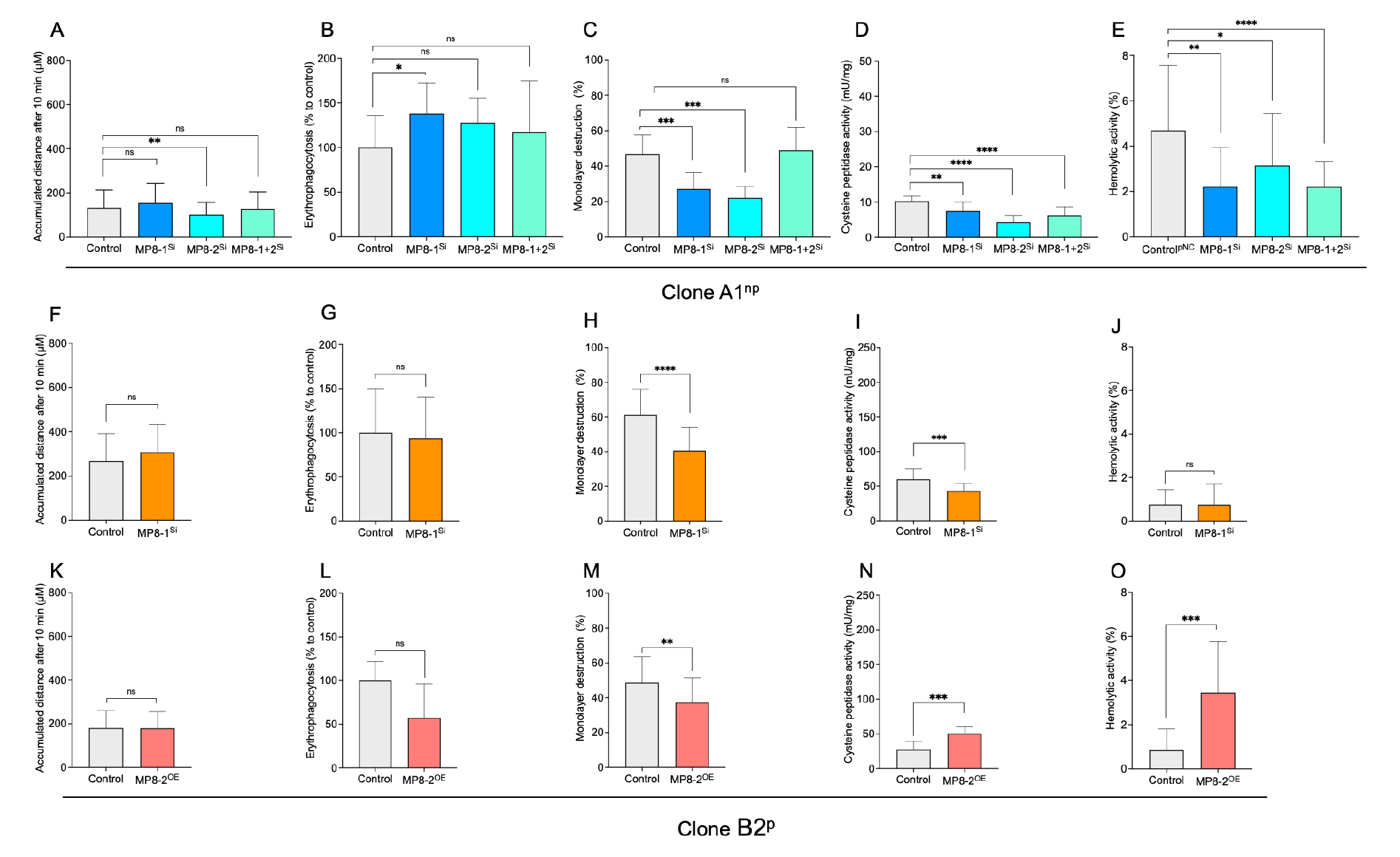
Determination of motility (A, F, K), erythrophagocytosis (B, G, L), cytopathic activity (C, H, M), cysteine peptidase activity (D, I, N), and hemolytic activity (E, J, O) of silencing (^Si^) and overexpression (^OE^) transfectants of clones A1^np^ (A1^np^MP8-1^Si^, A1^np^MP8-2^Si^, A1^np^MP8-1+2^Si^) and B2^p^ (B2^p^MP8-1^Si^, B2^p^MP8-2^OE^). A, F, K. To determine motility, the accumulated distance (µm) was measured after 10 min. For each transfectant/control 60 amoebae were analysed. B, G, L. To determine erythrophagocytosis, trophozoites (2 x 10^5^) and erythrocytes (2 x 10^8^) were incubated for 30 min at 37°C, non-phagocytosed erythrocytes were lysed, then the amoebae were lysed in 1% Triton-X-100 and absorbance was measured at 405 nm. The mean value of the controls was defined as 100% and the measured OD_405 nm_ values of the samples were related to it (6-9 biological replicates per transfectant/control). C, H, M. To determine cytopathic activity, HepG2 cells (1 x 10^5^) were seeded in 24 well plates, cultured for 48 h and stained with BCECF. Subsequently, 1 x 10^5^ trophozoites were added to the cells in 500 µl DMEM medium and incubated at 37° for 1 h. Afterwards, the cells were lysed, centrifuged and the supernatant was measured at 485 nm absorbance and 535 emission. The negative control was set at 100%. Experiments were performed at least 3 times in triplicate. D, I, N. Cysteine peptidase activity was determined using Z-Arg-Arg-pNA as substrate. Experiments were performed at least 9 times in duplicate. E, J, O. For hemolytic activity, 1.25 x 10^5^ trophozoites were mixed with 2.5 x 10^8^ erythrocytes in 1 ml of PBS and incubated at 37°C for 1 h. After incubation, the cells were sedimented, and the hemoglobin released in the supernatant was measured at 530 nm. Separately incubated erythrocytes and trophozoites were used as negative controls. To determine 100% hemoglobin release, 2.5 x 10^8^ erythrocytes were lysed in 1 ml of water. Experiments were performed at least 3 times in quadruplicate. ns: not significant, \**p*<0.05, \*\**p*<0.01, \*\*\**p*<0.001, \*\*\*\**p*<0.0001 (Mann-Whitney *U* test).

Substrate gel electrophoresis confirmed the results of the activity measurement. The overall decrease in activity of the A1^np^MP8-1^Si^ transfectant compared to the control is too small to be detected in a substrate gel. However, for the A1^np^MP8-2^Si^ transfectant, though, there is a clear decrease in the intensity of the EhCP-A1 and EhCPA-7 bands, and for the A1^np^MP8-1+2^Si^ transfectant, all bands are decreased in intensity. This is also very clearly confirmed for the B2^p^MP-1^Si^ transfectants. In particular, the activity bands of EhCP-A7 are clearly reduced in intensity (Fig 5).

**Fig 5.**
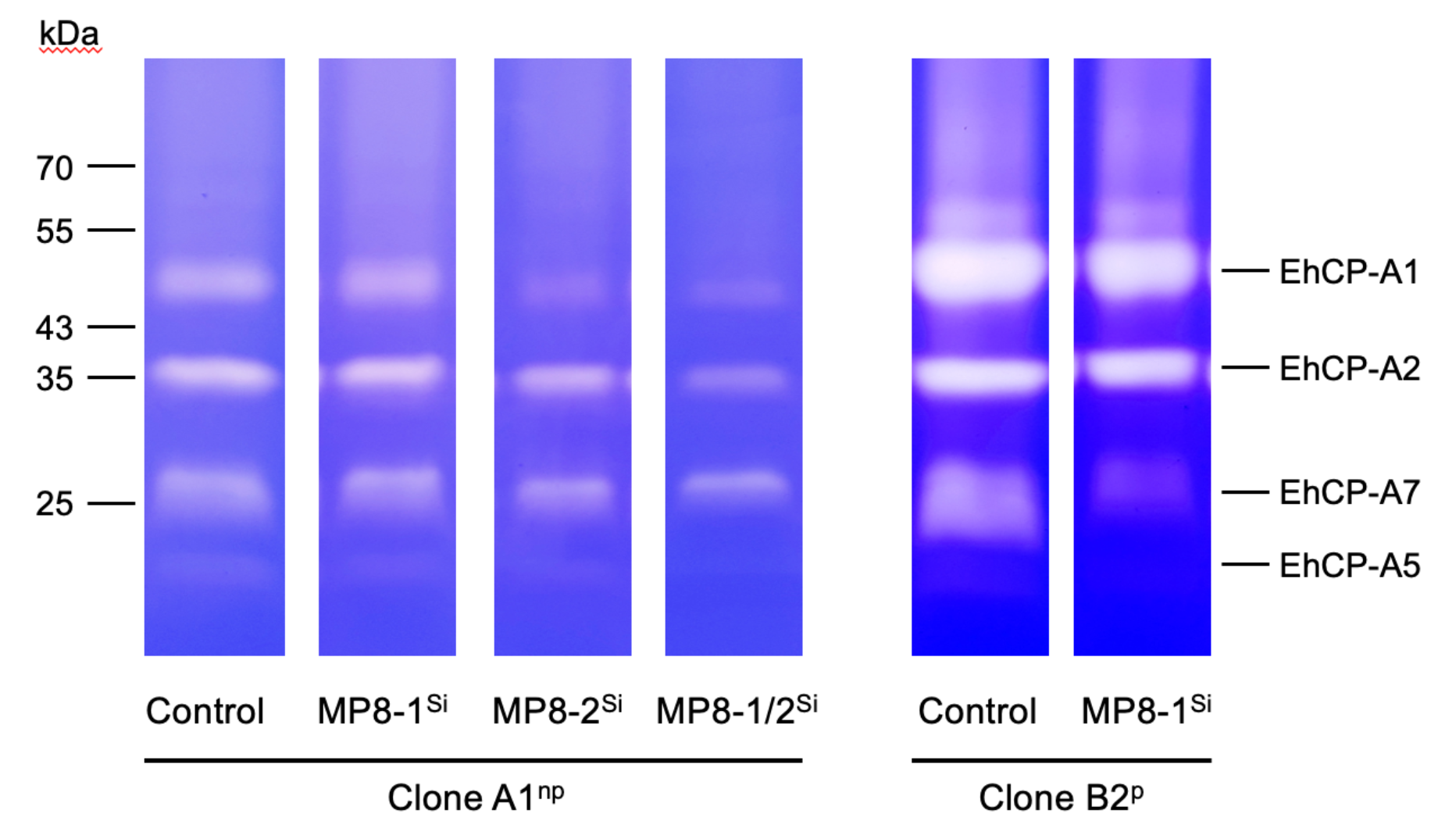
Determination of cysteine peptidase activity using substrate gel electrophoresis. 4 µg of amoeba extracts from controls (A1^np^/B2^p^) and transfectants (A1^np^MP8-1^Si^, A1^np^MP8-2^Si^, A1^np^MP8-1+2^Si^, B2^p^MP8-1^Si^) were separated on SDS-Page co-polymerized with gelatine. To visualize the cysteine peptidase activity, the gels were stained with Coomassie Blue.

Hemolytic activity is significantly reduced in all A1^np^ silencing transfectants (A1^np^EhMP8-1^Si^, *p*=0.0011; A1^np^EhMP8-2^Si^, *p*=0.0425; A1^np^EhMP8-1+2, *p*<0.0001). No effect was observed in B2^p^EhMP8-1^Si^ transfectants. However, B2^p^ trophozoites already have very low hemolytic activity (Fig 4E and 4J). Overexpression of the *ehmp8-2* gene in B2^p^ trophozoites led to a significant increase in hemolytic activity (B2^p^MP8-2^OE^, *p*=0.001) (Fig 4O).

### Silencing of *ehhp127* in pathogenic B2^p^ trophozoites and overexpression of *ehhp127* in non-pathogenic A1^np^ trophozoites

It has been shown that *ehhp127* is almost exclusively expressed in clone B2^p^ (reads: B2^p^-2314.73 vs A1^np^ 12.01) [4]. In contrast to the silencing of *ehmp8-1* and *ehmp8-2* genes, the silencing of *ehhp127* in B2^p^ trophozoites (B2^p^HP127^Si^) had no effect on the expression of other genes (Fig 6A, S5 Table). However, this is different when *ehhp127* was overexpressed in clone A1^np^ (S6 Table). In A1^np^HP127^OE^ transfectants, *ehhp127* was overexpressed approximately 5500-fold. Overall, 17 genes were significantly upregulated more than 3-fold and 40 genes were significantly upregulated more than 2- 3-fold significantly (padj<0.05) (Fig 6 B, S6 Table). However, when we compare the expression profile of the control transfectant A1^np^pNC with that of non-transfected A1^np^ trophozoites, we observed 242- and 7-fold repression of the 6 most differentially expressed genes between, whereas in A1^np^HP127^OE^ transfectants, the expression level returns to that of wild-type A1^np^ [4]. All 6 genes encode for hypothetical proteins (S6 Table). Why mock transfection had this effect is unknown.

**Fig 6.**
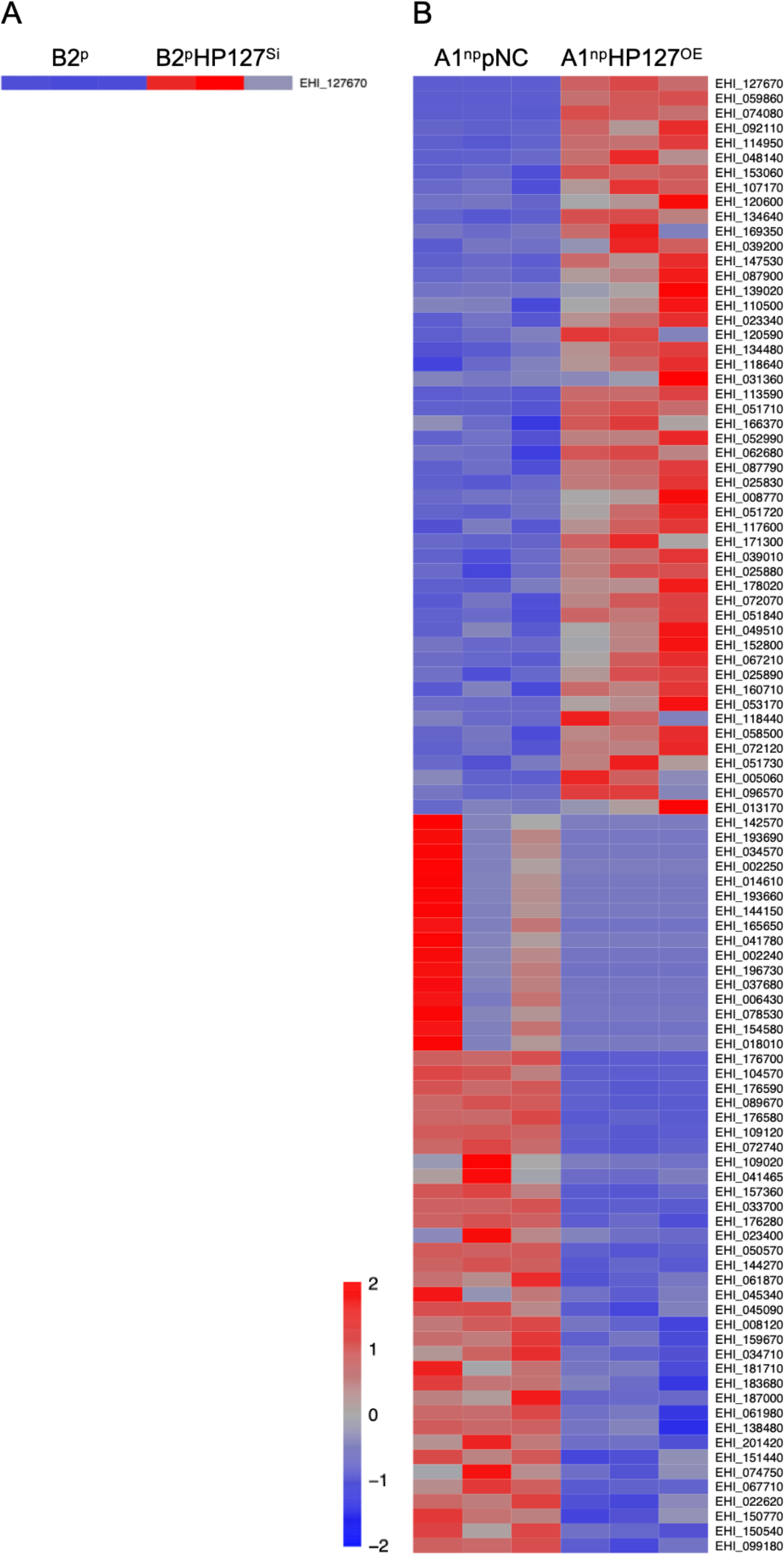
Heatmap of genes significantly differentially expressed (>1.8 fold, padj<0.05) in the different *E. histolytica* transfectants silencing or overexpressing *ehhp127* compared to the corresponding controls. A maximum of 50 up-and downregulated genes with the highest fold change are shown. A. B2^p^ versus B2^p^HP127^Si^, B. A1^np^pNC versus A1^np^HP127^OE^, C. A1^np^ versus A1^np^MP8-1+2^Si^, D. B2^p^ versus B2^p^MP8-1^Si^, E. B2^p^pNC versus B2^p^MP8-2^OE^.

Only one of the additional differentially expressed genes (*ehi_062680*), encoding a hypothetical protein, was also found to be significantly differentially expressed between wild-type A1^np^ and B2^p^ trophozoites. This gene was 20-fold more highly expressed in B2^p^ than in A1^np^ trophozoites [4]. The increase in expression from A1^np^pNC to A1^np^HP127^OE^ was 2.5-fold (S6 Table). However, transfection with the control plasmid pNC also appears to affect expression. The expression of a number of genes was up-or down-regulated compared to wild-type A1^np^, but also compared to A1^np^Eh127^OE^ transfectants (S6 Table). In total, the expression of 86 genes was significantly downregulated. Of these, 43 genes are downregulated >3-fold and another 43 up to 2-3-fold (S6 Table). Again, we see an effect of the mock transfection. The expression of 18 genes that were normally not or only weakly expressed in A1^np^ wild-type trophozoites occurs in A1^np^pNC. However, the expression of the A1^np^HP127^OE^ transfectants has returned to the low levels seen in A1^np^ trophozoites. The difference in expression here was between 4200- and 6.7-fold. Except for one *aig1* gene and one gene encoding a putative DNA polymerase, these are all genes encoding hypothetical proteins (Table 1). For all other genes that were significantly downregulated in A1^np^HP127^OE^ transfectants, the expression in A1^np^pNC control transfectants correlated with that in A1^np^ wild-type trophozoites [4]. It is therefore likely that this is a specific effect of *ehhp127* overexpression. It is striking that, overall, the expression of 11 *aig1* genes was significantly downregulated (Table 1). Furthermore, the expression of the genes encoding cysteine peptidases *ehcp-a4* and *ehcp-a6*, genes encoding heat shock proteins (HSP101, DNAJ family protein, HSP70) and genes encoding antioxidants (peroxiredoxin, iron-sulfur flavoprotein) was downregulated in A1^np^HP127^OE^ transfectants (S6 Table).

For nine genes, the expression profile of A1^np^pNC transfectants compared to A1^np^HP127^OE^ transfectants was comparable to that of A1^np^ to B2^p^ trophozoites. In both A1^np^HP127^OE^ transfectants and B2^p^ trophozoites, these genes are less expressed compared to A1^np^ control transfectants and A1^np^ trophozoites, respectively. These nine genes encode for heat shock proteins (EHI_034710, EHI_022620), AIG family proteins (EHI_126560, EHI_126550), DNA mismatch repair protein Msh2 (EHI_123830), splicing factor 3B subunit 1 (EHI_049170) and three hypothetical proteins (EHI_005657, EHI_075640, EHI_075690) (S6 Table).

### Phenotypic characteristics of EhHP127 transfectants

As with the EhMP8 transfectants, various phenotypic characteristics were also analyzed for the EhHP127 overexpressing and silencing transfectants (A1^np^HP127^OE^, B2^p^HP127^Si^).

Silencing and overexpression of *ehhp127* had less overall impact and less significant effects on the *E. histolytica* phenotype. For the silencing transfectants, this was not surprising since no other genes except *ehhp127* itself was affected in its expression. What is surprising, however, is that although the expression profile for a number of genes was regulated during overexpression, this only affected motility. This was also highly significantly affected in silencing transfectants. While overexpression of *ehhp127* in A1^np^ (A1^np^HP127^OE^) increases the distance traveled from 261±25 µm to 389±84 µm after 10 min (*p*<0.0001) (Fig 7A), silencing of *ehhp127* in B2^p^ (B2^p^HP127^Si^) led to a decrease in the distance traveled from 449±141 µm/10 min to 270±38µm/10 min (*p*<0.0001) (Fig 7F). Erythrophagocytosis was not affected in the *ehhp127* transfectants A1^np^HP127^OE^ and B2^p^HP127^Si^ (Fig 7B and 7G). However, as with all metallopeptidase transfectants, we observed that silencing of *ehhp127* reduced cytopathic activity. While 74±8.6 % of the monolayer was destroyed by the control (B2^p^), silencing of the *ehhp127* gene (B2^p^HP128^Si^) reduces the destruction to 51±10 % (*p*<0.0001) (Fig 7H). As previously described [5], overexpression had no significant effect on cysteine peptidase activity, whereas silencing of *ehhp127* results in a significant reduction of cysteine peptidase activity (190±49 mU/mg vs 123±23 mU/mg, *p*=0.0317) (Fig 7I). With respect to hemolytic activity, only a significant reduction in B2^p^ (2.6±1.2 % vs 1.7±1 %, *p*=0.0328) was observed after *ehhp127* silencing (Fig 7J).

**Fig 7.**
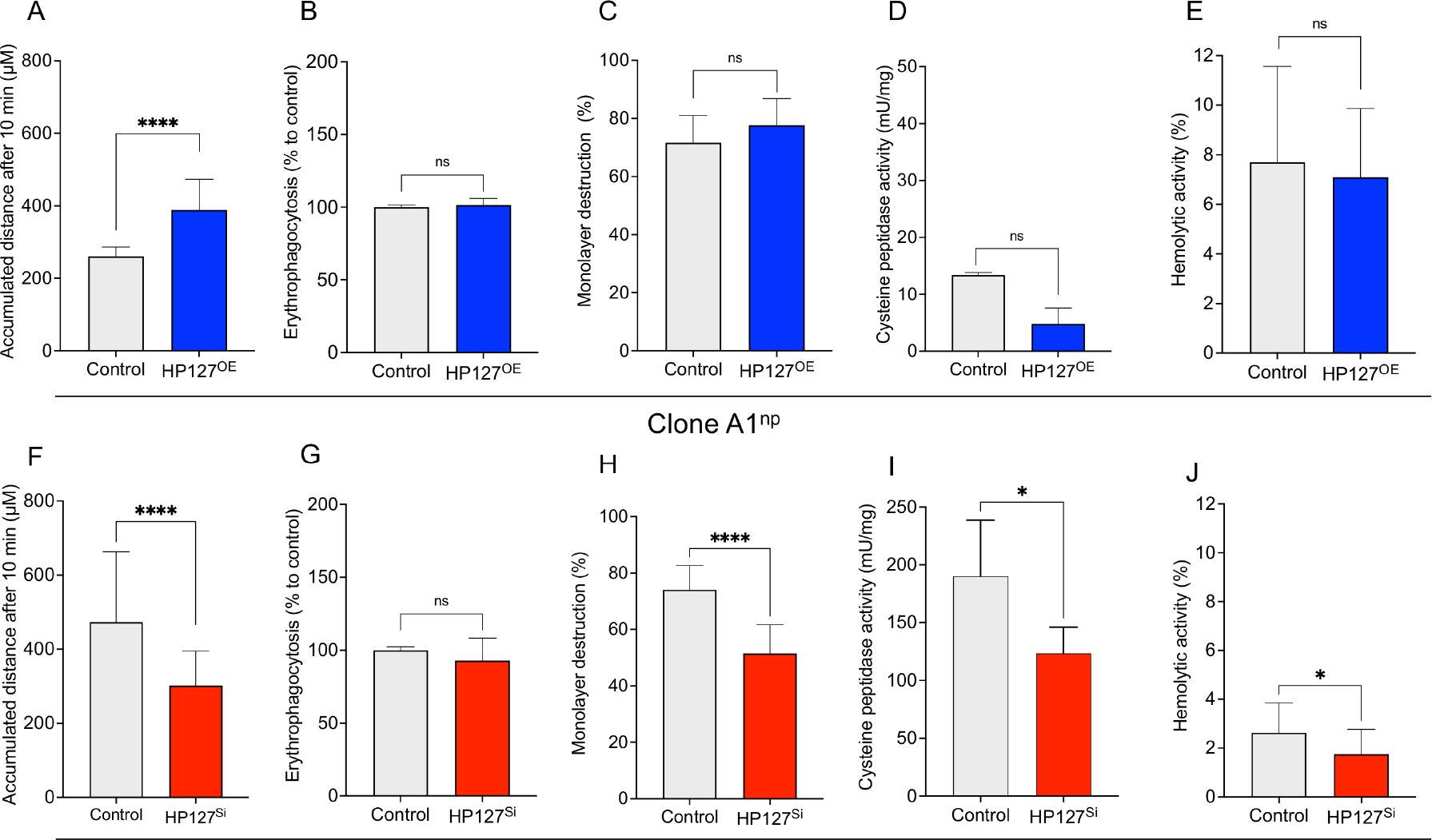
Determination of motility (A, F), erythrophagocytosis (B, G), cytopathic activity (C, H), cysteine peptidase activity (D, I), and hemolytic activity (E, J) of EhHP127 transfectants (overexpression transfectant: A1^np^HP127^OE^, silencing transfectant: B2^p^HP127^Si^). A, F. To determine motility, the accumulated distance (µm) was measured after 10 min. For each transfectant/control 30 amoebae were analyzed). B, G. To determine erythrophagocytosis, trophozoites (2 x1 0^5^) and erythrocytes (2 x 10_8_) were incubated for 30 min at 37°C, non-phagocytosed erythrocytes were lysed, then the amoebae were lysed in 1 % Triton-X-100 and absorbance was measured at 405 nm. The mean value of the controls was defined as 100%, and the measured OD_405 nm_ values of the samples were related to 100% (5-12 biological replicates per transfectant/control). C, H. To determine cytopathic activity, HepG2 cells (1 x 10^5^) were seeded in 24 well plates, cultured for 48 h and stained with BCECF. Subsequently, 1×10^5^ trophozoites were added to the cells in 500 µl DMEM medium and incubated for 1 h at 37°C. Afterwards, the cells were lysed, centrifuged and the supernatant was measured at 485 nm absorbance and 535 nm emission. The negative control was set at 100 %. Experiments were performed at least 3 times in duplicate. D, I. Cysteine peptidase activity was determined using Z-Arg-Arg-pNA as substrate. The experiments were performed in duplicate at least 3 (D) and 5 (I) times. E, J. For the determination of hemolytic activity 1.25 x 10^5^ trophozoites were mixed with 2.5 x 10^8^ erythrocytes in 1 ml of PBS and incubated for 37°C at 1 h. After incubation, the cells were sedimented and the hemoglobin released in the supernatant was measured at 530 nm. Separately incubated erythrocytes and trophozoites were used as negative controls. To determine 100% hemoglobin release, 2.5 x 10^8^ erythrocytes were lysed in 1 ml of water. The experiment was performed 2 times in duplicate (E) and 7 times in triplicate (J). ns: not significant, \**p*<0.05, \*\*\*\**p*<0.0001 (Mann-Whitney *U* test).

### Comparison of EhMP8-1 and EhMP8-2 with leishmanolysins and invadolysins of different organisms

Comparing the amino acid sequences of EhMP8-1 and EhMP8-2 with leishmanolysin-like peptidases from different organisms, the percent identity to those of the liver fluke *Clonorchis sinensis*, the avian protozoan parasite *Histomonas meleagridis*, and *T. vaginalis* ranges from 16 % to 20 %. The identity to leishmanolysin from *L. major* is 22 % to 23 %. The highest identity (24 % to 28 %) is found with various metazoan invadolysins such as those from the hookworm *Ancylostoma caninum*, the whipworm *Trichuris suis*, *Drosophila melanogaster*, *Mus musculus*, *Macaca thibetana*, and humans, as well as the soil-dwelling amoeba *Dictyostelium discoideum* (S7 Table).

### Localization of EhMP8-1, EhMP8-2 and EhHP127

To localize the two metallopeptidases and the hypothetical protein EhHP127, amoebae were transiently transfected with plasmids, allowing translation of the respective protein fused to a cMyc-tag at the C-terminus.

In the pNCMP8-1^Myc^ and pNCMP8-2^Myc^ transfectants, both metallopeptidases were detected in vesicle-like structures within the trophozoites. They mainly localized to vesicles located in the endoplasm of trophozoites (Fig 8A-8F). Neither EhMP8-1 nor EhMP8-2 were detected on the cell surface of trophozoites.

**Fig 8.**
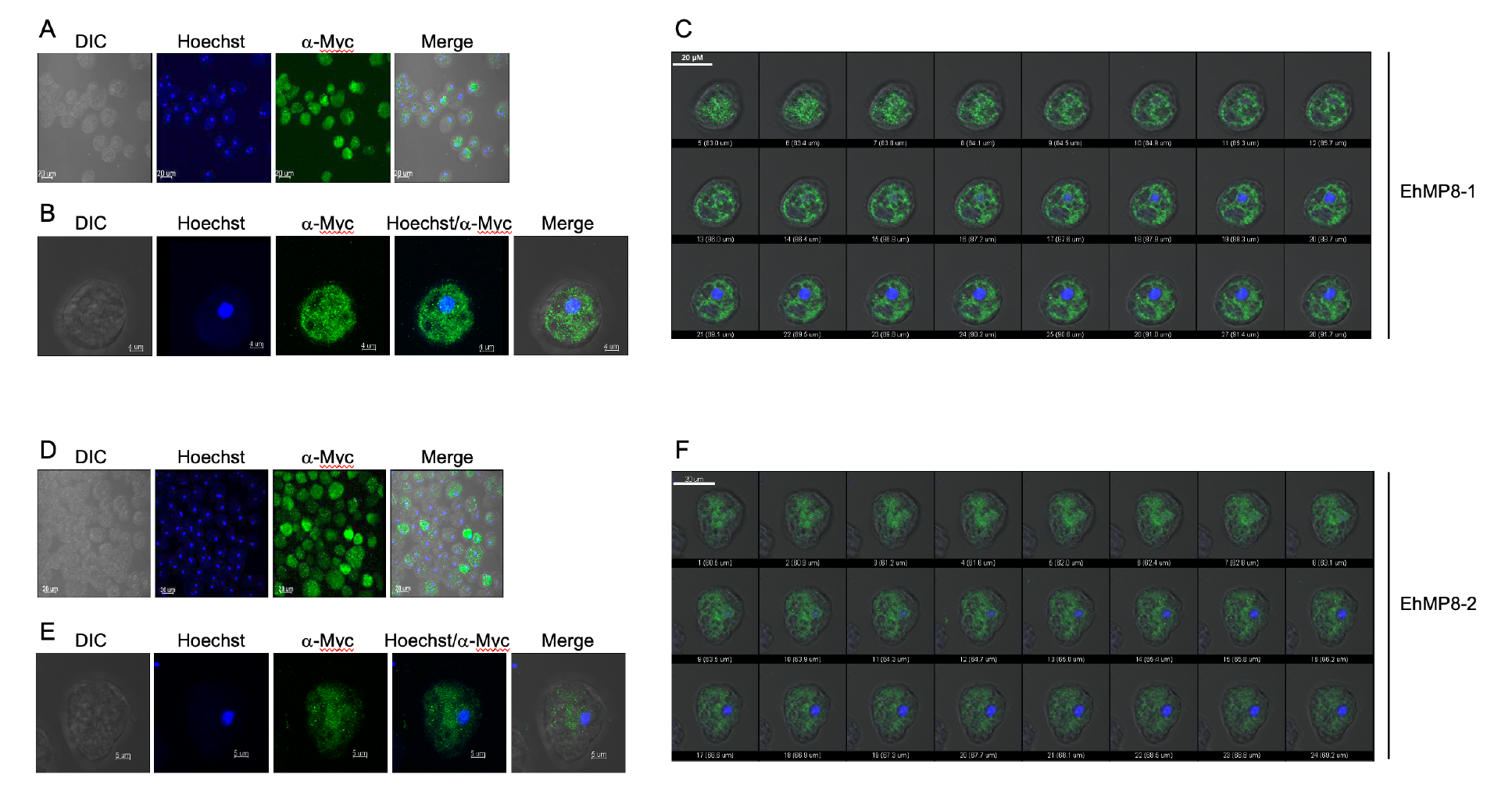
Microscopic analysis of A1^np^MP8-1^Myc^ and A1^np^MP8-2^Myc^ transfectants for localization of the metallopeptidases EhMP8-1 and EhMP8-2. The A1^np^ trophozoites were transfected with the expression plasmids pNCMP8-1^Myc^ or with pNCMP8-2^Myc^, which allowed the production of a metallopeptidase-myc fusion protein, and the myc-tag could be stained with a specific anti-myc antibody. For the immunofluorescent analysis, trophozoites were fixed with PFA and permeabilized with saponin. MP8-1-Myc and MP8-2 fusion proteins were stained with ⍺-c-myc primary antibody (1:100) and ⍺-mouse Alexa Fluor 488 (1:400, green). A, D. Overview of trophozoites of the A1^np^MP8-1^Myc^ (A) and A1^np^MP8-2^Myc^ (D) transfectant. B, E. Single trophozoite of the A1^np^MP8-1^Myc^ (B) and A1^np^MP8-2^Myc^ transfectant (E). Shown are transmitted light images (DIC), staining of nuclei with Hoechst dye, staining of EhMP8-1 and EhMP8-2 with anti-Myc antibody, and an overlay of the images (merge). C, F. Single images of confocal micrographs of a trophozoite of the A1^np^MP8-1^Myc^ (C) and A1^np^MP8-2^Myc^ (F) transfectant. The single images show 0.38 μm thick sections through the trophozoites. A total of 42 images were obtained, and only every second image is shown here.

In the B2^p^EhHP127^Myc^ transfectants, EhHP127^Myc^ also localized to trophozoite vesicles. A uniform distribution within the trophozoites was observed (Fig 9A-9C). In a Western blot analysis, EhHP127 was detected only in the insoluble pellet fraction and not in the soluble fraction (Fig 9D).

**Fig 9.**
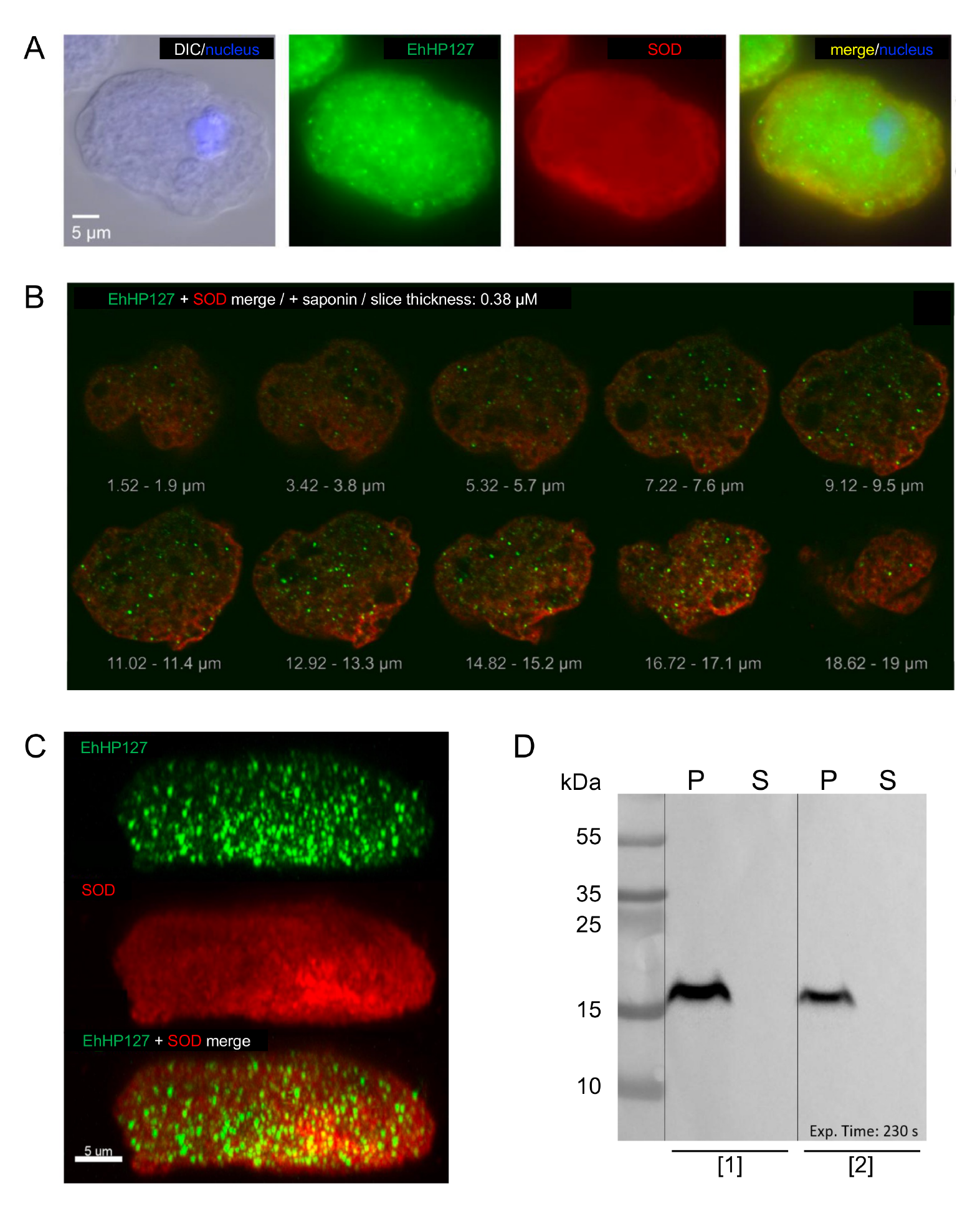
Localization of the hypothetical protein EhHP127. B2^p^ trophozoites were transfected with the expression plasmid pNCHP127^Myc^, which allowed the production of an EhHP127-Myc fusion protein, and the myc-tag was stained with a specific anti-myc antibody. A-C. For the immunofluorescent analysis, trophozoites were fixed with PFA and permeabilized with saponin. EhHP127-Myc fusion protein was stained with ⍺-c-myc primary antibody (1:100) and ⍺-mouse Alexa Fluor 488 (1:400, green). For co-localization an antibody against the cytoplasmically localized superoxide dismutase (SOD) and ⍺-rabbit Alexa Fluor 594 (1:400, red) was used. Nuclei were stained with Hoechst dye (blue). A. Single trophozoite from the B2^np^HP127^Myc^ transfectant. B. Confocal images of a trophozoite of the B2^p^Eh127Myc transfectant. Series of 10 sections with a distance of 1.9 μm from top to bottom, covering a major part of the cell. C. Side view of the whole cell with the three modes green, red and merge, for the detection of EHI_127670 and SOD both single and merged. For the side view, a 3D-model was constructed out of 55 single slices using Imaris. D. Western blot of a 13 % Tricine-SDS-PAGE using an ⍺-c-myc primary antibody (1:1000) and ⍺-mouse-HRP secondary antibody (1:2000). For the Western blot, B2^p^ trophozoites were transfected twice independently ([1], [2]) and the pellet fraction (P) and the soluble fraction (S) were prepared from each of the two transfectants.

## Discussion

Comparative transcriptome analysis of pathogenic and non-pathogenic clones, all derived from the *E. histolytica* isolate HM-1:IMSS, identified, among others, two differentially expressed genes whose silencing or overexpression affects the virulence phenotype of the amoebae. Overexpression of the gene *ehmp8-2*, encoding the metallopeptidase EhMP8-2, was shown to decrease the virulence of the pathogenic clone B2^p^, whereas overexpression of the gene *ehhp127*, encoding a hypothetical protein, increased the virulence of the non-pathogenic clone A1^np^ [4, 5]. To better understand the function of these two proteins, corresponding silencing or overexpression transfectants were generated and their phenotypes analyzed.

As described above, *E. histolytica* has two metallopeptidases, EhMP8-1 and EhMP8-2, both belonging to the M8 peptidase family (leishmanolysin-/gp63-like). In the protozoan parasite *Leishmania*, leishmanolysin is essential for its virulence [11]. This peptidase is the most abundant protein on the cell surface during the promastigote stage of the parasite, and is attached to the membrane by a glycosylphosphatidylinositol anchor. This peptidase acts as a ligand involved in the direct interaction of promastigotes with host macrophage receptors, interaction with the complement cascade, and is thought to facilitate the migration of the organism through the extracellular matrix of the host [12-15]. For *Leishmania*, but also for *Trypanosoma*, an involvement in the attachment to the midgut of the insect vector has also been described [16, 17]. Furthermore, *Trypanosoma* leishmanolysin is involved in the differentiation from the bloodstream form to the procyclic form [18]. In *L. braziliensis*, 29 leishmanolysin genes have been identified, clustered into three different subgroups [19]. In *Trichomonas,* 42 leishmanolysin-like family members have been annotated. An M8 family member was reported to play a role in *T. vaginalis* infection and a surface proteomics approach identified 16 leishmanolysin-like proteins [20, 21]. However, leishmanolysin-like peptidases are not unique to parasitic protozoa, but are conserved in many different organisms, and the number of gene copies can vary substantially. In contrast to *Leishmania*, trypanosomes, or trichomonads, as described above *E. histolytica* has only two members of the M8 family. In this context, it is even more surprising that non-pathogenic A1^np^ trophozoites express both genes, while pathogenic B2^p^ trophozoites still have two intact genes but no longer express the *ehmp8-2* gene [4]. Leishmanolysin-like proteins have also been described in metazoans, where they are called invadolysins [22]. In *Drosophila melanogaster* or in humans, only one gene encoding an invadolysin is found, although there are four splice variants in humans [23]. The identity of EhMP8-1 and EhMP8-2 with leishmanolysins and invadolysins of other organisms is relatively low, namely 16%-28%. Interestingly, the metallopeptidases of *E. histolytica* show the highest identity to several invadolysins of metazoans (S7 Table).

Invadolysin has been shown to be essential for larval survival in *Drosophila*. Mutants lacking the gene die in the late larval stages and produce abnormal mitotic phenotypes in neuroblasts [22]. Other phenotypes associated with invadolysin mutants include increased levels of nuclear envelope proteins and impaired germ cell migration in embryos [22]. In human cells, invadolysin can be detected in the nucleus. However, in both *Drosophila* and human cells, it is localized in the cytoplasm, where it is concentrated in unusual ring-like structures [22]. A later study showed that the ring-shaped subcellular localization of invadolysin surrounds lipid droplets and that invadolysin is located on the surface of these lipid droplets [23].

We detected both EhMP8-1 and EhMP8-2 in vesicles in the amoebae (Figure 8A-F). In contrast to Teixera and colleagues [7], who sporadically detected EhMP8-1 on the cell surface of amoebae, we could not verify this localization. It is not yet clear what kind of vesicles these are, or whether they could be some kind of lipid droplets. However, to date, no lipid droplets have been detected in *E. histolytica* using a specific lipid stain. In stationary macrophages, invadolysin was localized in lipid droplets in the cytoplasm, but in actively migrating macrophages it was highly concentrated at the leading edge of the cell in actively migrating macrophages, which also provides a link to cell migration [22]. However, the relationship between lipid droplets and cell migration is not fully understood. It has also been shown that increased numbers of lipid droplets have been observed in neoplastic colon cancer cells [24], so it is possible that these organelles may play a role in the transformation and spread of cancer cells. It has also been shown that inhibition of intracellular membrane trafficking blocks extracellular matrix degradation by invadopodia [25], and since endocytosis is associated with cell migration [26, 27], it is possible that lipid droplet-associated invadolysin affects cell migration via the invadopodia [23]. This parallels studies on EhMP8-1. Hasan and colleagues showed that silencing of *ehmp8-1* (*ehmsp-1*) renders trophozoites hyperadherent and less motile. They also showed that silencing of *ehmp8-1* results in parasites that are unable to form specialized, dot-polymerized actin structures (F-actin) upon interaction with human fibronectin. These short-lived F-actin structures resemble those of mammalian cell invadopodia [28]. In contrast to the study by Hasan and colleagues, we did not observe any effect of *ehmp8-1* silencing on motility (Figure 4A and 4F). However, we showed that both silencing and overexpression of *ehmp8-2* and its homologous gene *ehmp8-1* have the strongest effects on cysteine peptidase and hemolytic activity. Silencing leads to a decrease in the corresponding activities, whereas overexpression leads to an increase (Fig 4D, 4E, 4I, 4J, 4N, 4O). The ability to disrupt a monolayer is also significantly affected, with reduced cytopathic activity in both overexpressing and silencing transfectants (Fig 4C, 4H, 4M).

To determine whether the altered phenotypes were indeed due to silencing or overexpression of the corresponding *ehmp8* genes, the expression profiles of the different transfectants were analyzed by RNAseq. It was striking that silencing *ehmp8-2*, but also the homologous gene *ehmp8-1* of both genes affected the expression of several hundred genes in the transfectants. It is not known why silencing has such a strong effect on the expression of many other genes. As many of these genes encode proteins of unknown function, only limited statements can be made about the metabolic pathways that might be affected. Remarkably, silencing of metallopeptidase genes led to upregulation of genes encoding antioxidants including genes encoding iron-sulfur flavoproteins. In a comparative study between the pathogenic isolate HM-1:IMSS and the non-pathogenic isolate UG10 (derived from HM-1:IMSS), several iron flavoproteins were also identified as differentially upregulated in UG10 [29]. In addition, the expression of a number of *aig1* genes is downregulated in the different EhMP8 silencing transfectants (Table 1). An influence on the expression of *aig* genes is also seen in the EhHP127^OE^ overexpression transfectants. In these, the expression of ten *aig1* genes is significantly downregulated (Table 1). AIG1 proteins were first described in the context of plant immune defense [30]. The function of these molecules in *E. histolytica* has not yet been elucidated. More than 45 *aig1* genes have been described [2]. The proteins encoded by them show structural similarities to the GTPases of immunity-associated protein (GIMAPS)/immune-associated nucleotide-binding protein (IAN) family of AIG1-like GTPases, which are conserved between vertebrates and angiosperms plants [31]. Comparison of a pathogenic with a non-pathogenic HM-1:IMSS cell line, from which the clones studied here were derived, showed that of 34 *aig1* genes examined, 18 genes were more highly expressed in the pathogenic cell line B and only one gene was less highly expressed than in the non-pathogenic cell line A [2]. Some of the AIG1 proteins were found to have specific functions. For example, it was shown that overexpression of the *aig1* gene EHI_176590 in strain HM-1:IMSS cl6 resulted in increased formation of cell surface protrusions and increased adhesion of trophozoites to human erythrocytes [32]. Overexpression of the *aig1* gene EHI_180390 in trophozoites of the non-pathogenic strain UG10 resulted in increased virulence. Furthermore, co-culture of pathogenic HM-1:IMSS with *Escherichia coli* reduced virulence *in vitro* and downregulated *aig1* gene expression. In contrast, the virulence of strain UG10 co-cultured with *E. coli* was increased and *aig1* expression was upregulated [29]. It has also been shown that an *aig1* gene is upregulated in its expression in the presence of H_2_O_2_ [33]. In amoebae resistant to 12 µM metronidazole, 10 *aig1* genes were differentially expressed [34]. Similarly, differential expression was detected during *E. histolytica* invasion of the mouse intestine [35], and *aig1* genes were downregulated in trophozoites isolated from an amoebic liver abscess [36]. Three members of the *aig* family have also been identified as cyst-specific genes [37].

However, particularly in the EhHP127^OE^ overexpression transfectants, it is apparent that although the expression of 10 *aig1* genes is downregulated, and the expression of a variety of other genes is altered, this has no significant effect on most of the phenotypes examined here. Overexpression of *ehhp127* only results in a significant increase in motility. In contrast, silencing of the *ehhp127*, which does not alter the expression of other genes, significantly reduces motility. It is therefore very likely that EhHP127 affects the mobility of amoebae. However, the mechanism is not known. The molecule has no homologue in the animal kingdom and no specific domains have been identified, so it is not possible to hypothesize about its function.

Silencing of *ehhp127* also results in reduced cytopathic, cysteine peptidase and hemolytic activity. That silencing of *ehhp127* has an effect on cysteine peptidase activity was previously shown by Matthiesen and colleagues and was confirmed here [5].

Interestingly, some cysteine peptidases have been identified in trophozoite vesicles [38-44], such as EhHP127, but also EhMP8-1 and EhMP8-2, for which a direct or indirect effect on cysteine peptidase activity could be demonstrated using the corresponding transfectants.

In conclusion, the overexpressing and silencing transfectants studied here do not provide a clear indication of the function of the metallopeptidases EhMP8-1 and EhMP8-2 and the hypothetical protein EhHP127. However, it was surprising to observe that both silencing and overexpression of metallopeptidase genes have a strong effect on the expression of a large number of other genes. This suggests that they play an important role in regulating gene expression. In a previous study, the motility of B2^p^ trophozoites was shown to be significantly higher than that of A1^np^ trophozoites [4]. Furthermore, the overexpression and silencing transfectantion experiments performed here clearly show that EhHP127 appears to be a protein that affects amoeba motility in a previously unknown way. Whether there is also a link between motility and pathogenicity cannot be conclusively determined, but these results imply that enhanced motility is a phenotype that is causally related to virulence.

## Material and Methods

### *E. histolytica* cell culture and generation of transfectants

Cultivation of *E. histolytica* trophozoites was performed under microaerophilic and axenic conditions at 37°C in plastic culture flasks (Corning, Kaiserslautern, Germany) using TYI-S-33 medium [45]. Non-pathogenic *E. histolytica* clone A1^np^ and pathogenic *E. histolytica* clone B2^p^ were derived from the cell lines HM-1:IMSS-A and HM-1:IMSS-B [4].

Overexpressing and silencing transfectants were generated as described [4, 5]. In summary, for overexpression of *ehmp8-2* (*ehi_042870*) in clone B2^p^ and *ehhp127* (*ehi_127670*) in clone A1^np^, trophozoites were transfected with the expression plasmid pNC containing the gene of interest under control of the *E. histolytica* lectin promoter (B2^p^MP8-2^OE^, A1^np^HP127^OE^). As a control B2^p^ or A1^np^ trophozoites were transfected with the plasmid pNC [4]. Overexpressing transfectants were cultured in TY-I-S-33 medium containing 20 µg/ml G-418 (Gibco, Thermo Fisher Scientific, Schwerte, Germany). For silencing of *ehmp8-1* (*ehi_200230*) or *ehmp8-2* in clone A1^np^, trophozoites were transfected with the silencing plasmid pSiA containing *ehmp8-1* (267 bp) or *ehmp8-2* (1989 bp), respectively, in frame with the trigger region of *ehi_169670* (A1^np^MP8-1^Si^, A1^np^MP8-2^Si^). For silencing of *ehmp8-1* or *ehhp127* in clone B2^p^, trophozoites were transfected with the silencing plasmid pSiB containing *ehmp8-1* (268 bp) or *ehhp127* (917 bp) in frame with the trigger region of *ehi_074080* (B2^p^MP8-1^Si^) [5]. The silencing transfectants were grown in TY-I-S-33 medium containing 20 µg/ml G-418 for 3 weeks. After cloning of the transfectants by limited dilution, the cells were cultured without selection for at least 4 months until complete loss of the plasmid. Plasmid loss was proven by culturing the amoebae in the presence of 20 µg/ml G418 [5]. A1^np^ and B2^p^ trophozoites were used as controls. Overexpression and silencing were confirmed by specific qRT-PCR (S8 Table).

For localization studies, *ehhp127* (*ehi_127670*) was expressed under its own promoter using a myc-tag containing expression vector (pNC^Myc^) in clone B2^p^. The pNC^Myc^ plasmid is based on the pNC expression plasmid. A myc-tag was integrated into the *Bam*HI restriction site via *Bam*HI/*Bgl*II. To generate the pNC^Myc^ vector, two complementary oligonucleotides encoding the cMyc-tag (Glu-Gln-Lys-Leu-Ile-Ser-Glu-Glu-Asp-Leu) and the restriction sites *Kpn*I, *Nhe*I, *Bam*HI, *Xho*I, and *Bgl*II at the 5’-end were hybridized (S8 Table). The resulting DNA fragment was then cloned into the pNC vector via the *Kpn*I restriction site at the 5’ end and the *Bgl*II restriction site at the 3’ end.

To generate the plasmid pNC_HP127^Myc^, a 500 bp long sequence upstream of the start ATG, presumably containing the promoter region, and the entire open reading frame of *ehi_127670* was amplified by PCR using a forward primer containing a *Kpn*I restriction site and the reverse primer containing an *Bam*HI restriction site (S8 Table). DNA from the clone B2^p^ was used as a template. The amplified insert was cloned into the pNC^Myc^ vector with *Kpn*I and *Bam*HI restriction sites.

Expression of *ehmp8-1* and *ehmp8-2* was also performed under their own promoter (pNCMP8-1^Myc^, pNCMP8-2^Myc^). For this purpose, a 500 bp long sequence upstream of start ATG, which presumably contains the promoter region, and the entire open reading frame of *ehmp8-1* (*ehi_200230)* and (*ehmp8-2) ehi_042870* was amplified by PCR using the forward primer containing a *Kpn*I restriction site and the reverse primer containing a *Bam*HI restriction site (S8 Table).

### RNA extraction and quantitative real-time PCR (qRT-PCR)

For RNA isolation, trophozoites were harvested, washed twice with sodium phosphate-buffered saline (NaPBS; 4°C; 6.7 mM NaHPO_4_, 3.3 mM NaH_2_PO_4_, 140 mM NaCl, pH 7.2) and lysed with Trizol reagent (QIAzol Lysis reagent, QIAgen, Hilden, Germany). Total RNA isolation and DNA digestion were performed using the Direct-zol^TM^ RNA MiniPrep kit (Zymo Research, Irvine, CA, USA). All steps were performed according to the manufacturer’s instructions. RNA concentration and purity were determined using the NanoDrop 2000 (Thermo Fisher Scientific, Schwerte, Germany). The cDNA required for qRT-PCR was synthesized from the isolated RNA using the SuperScriptIII First-Strand Synthesis System Kit (Invitrogen, Thermo Fisher Scientific, Dreieich, Germany), according to the manufacturer’s instructions.

Sense and antisense primers were designed for qRT-PCR experiments to amplify 80-120 bp fragments of the genes of interest (S8 Table). The Luna Universal qPCR Master Mix kit (New England Biolabs, Frankfurt, Germany) was used to perform qRT-PCR. 2-4 biological replicates were analyzed in duplicate each time. Relative concentrations in gene expression were calculated using the 2^-ΔΔCT^ method and Rotor Gene software (Rotor Gene 6, Corbett Research). A1^np^, B2^p^, and B2^p^pNC were used as calibrators (control), and were set to 1. Actin was used as a housekeeping gene for normalization.

### Next generation sequencing

The quality of purified RNA was assessed using the Agilent 2100® Bioanalyzer System (Agilent Technologies, Santa Clara, CA, United States) and the Agilent RNA 6000 Pico Reagents Kit (Agilent Technologies). Ribosomal RNA was removed using the QIAseq FastSelect RNA Removal Kit (Qiagen, Hilden, Germany) according to the manufacturer’s instructions. RNA from each sample was prepared for sequencing using the QIAseq Stranded mRNA Library Kit (Qiagen, Hilden, Germany) according to the manufacturer’s instructions. Normalized libraries were pooled and sequenced using a NextSeq 500/550 Mid OutputKit v2.5 (Illumina, San Diego, CA, USA) with 150 cycles (2 × 75 bp paired-end) on a NextSeq 550 platform, generating a depth of 5-16 million paired-end reads for each sample. Reads were trimmed and filtered using Trimmomatic [46], and reads were aligned to the *E. histolytica* transcriptome (AmoebaDB, Release 61, 15 Dec 2022) using RSEM [47] and Bowtie2 [48] software. Differential expression was tested using DEseq2 to normalize the raw data [49].

### Determination of motility

24 h before the experiment, 2.5×10^5^ trophozoites were seeded into a T25 culture flask (Sarstedt, Nümbrecht, Germany). For each experiment, 3 biological replicates were used and the speed of movement was determined for 20 amoebae each using an Evos FL Auto microscope (Thermo Fisher Scientific). A video was recorded for 10 min (one frame every 5 sec). Manual tracking of the amoebae was performed using ImageJ version 2.0.0-rc-43/1.51d with plugins for manual tracking and chemotaxis. Significance was determined using the Mann-Whitney *U* test.

### Erythrophagocytosis assay

Human erythrocytes (blood group 0+ donated from the blood bank of the University Medical Center Hamburg-Eppendorf) and trophozoites to be examined were washed twice with incomplete TY-I-S-33 medium (200 x *g*, 10 min, 4°C). Subsequently, 2 x 10^8^ erythrocytes and 2 x 10^5^ trophozoites (ratio 1000:1) were added to a 5 ml tube with a total volume of 400 μl incomplete TY-I-S-33 medium. After incubation at 37 °C for 30 min, 1 ml each of ddH_2_O was added twice and incubated for 1 min to lyse the non-phagocytized erythrocytes. The cells were then centrifuged at 400 x *g* for 4 min in a 15 ml tube and washed twice with NaPBS. The amoebae were then lysed to release the hemoglobin of the phagocytosed erythrocytes. For this purpose, 1 ml of 1% Triton-X 100 was added to the trophozoites. For photometric measurement, 200 μl of the lysed trophozoites were pipetted into a 96-well plate and measured at 405 nm. The mean value of each control was defined as 100 % and the measured OD values were referenced to this value. At least 5 biological replicates were performed. Significance (*p*- values) was determined using the Mann-Whitney *U* test.

### Determination of cytopathic activity (monolayer destruction)

To determine the cytopathic activity of *E. histolytica*, 1 x 10^5^ cells of the hepatocyte cell line HepG2 (Merck, Darmstadt, Germany) were seeded in the wells of a 24-well plate 48 h before the experiment and incubated at 37°C in 2 ml HepG2 cell culture medium (Advanced DMEM; Gibco, Thermo Fisher Scientific, Schwerte, Germany) supplemented with 10 % fetal calf serum (FCS; Capricorn, Hamburg, Germany) and 1x penicillin/streptomycin (Capricorn), with the medium changed once after 24 h. After 48 h, a confluent cell monolayer was formed. Trophozoites were seeded into a T12.5 culture flask 24 h prior to the experiment so that a confluent monolayer was formed the next day. At the beginning of the experiment, HepG2 cell culture medium was removed from the HepG2 cells, which were then washed with NaPBS. To stain the HepG2 cells, 200 μL NaPBS + 10 µM of the fluorescent dye BCECF, AM (2’,7’-bis-(2-carboxyethyl)-5-(and-6)-carboxyfluorescein, acetoxymethyl ester) was added. The cells were incubated at 37°C for 30 min. Afterwards, BCECF/NaPBS was removed, and the cells were washed twice with NaPBS (37°C). Then, 1 × 10^5^ trophozoites in 500 μl were added to the stained HepG2 cells. As a negative control, the stained HepG2 cells were incubated with 500 μl Advanced DMEM medium, while as a positive control, the stained HepG2 cells were lysed with 500 μl DMEM+1 % Triton X-100. The 24-well plate was then incubated at 37°C for 1 h. Subsequently, the plate was then placed on ice for 20 min to dissolve the trophozoites. After washing twice with NaPBS (4°C), 1 ml of 1 % Triton X-100 PBS was pipetted into each well to lyse the HepG2 cells, and the plate was incubated at 37°C for 30 min. 150 μl of the supernatant were pipetted into a black 96-well plate (Greiner, Frickenhausen, Germany), which was centrifuged at 1400 x *g* for 30 sec and measured with a fluorescence plate reader (GENios, TECAN, Fornax Technologies GmbH, Uetikon am See, Switzerland) at an absorbance of 485 nm and an emission of 535 nm. The negative control values were set to 100 %. The values of the samples were then related to the negative control. Finally, the result was subtracted from 100 % to determine the percentage of cells detached from the cell layer by the trophozoites. Experiments were performed at least 3 times in triplicate. Significance was determined using the Mann-Whitney *U* test.

### Determination of cysteine peptidase activity and substrate gel electrophoresis

Amoebae were seeded in T25 cell culture flasks 24 h prior to the start of the experiment to form a monolayer the next day. To prepare the amoeba extracts used for the determination of cysteine peptidase activity and substrate gel electrophoresis, the amoebae were lysed in liquid nitrogen over 3 freeze-thaw cycles and sedimented at 12,000 x *g* for 15 min at 4°C. The supernatant was used in the corresponding experiments. Protein concentration was measured using the Pierce™ BCA Protein Assay Kit (Thermo Fisher Scientific) according to the manufacturer’s instructions.

Cysteine peptidase activity was measured in a 96-well plate in a final volume of 200 µl (148 µl CP assay buffer (0.1 M KH_2_PO_4_, 2 mM EDTA, 1 mM DTT), 2 µl amoeba extract and 50 µl of the synthetic peptide Z-Arg-Arg-pNA (400 µM concentration; Bachem, Bubendorf, Switzerland) as substrate. One unit of enzymatic activity is defined as the amount of enzyme that catalyzes the formation of 1 mmol p-nitroaniline in 1 min. Experiments were performed at least 3 times in duplicate. Significance was determined by the Mann-Whitney *U* test.

Substrate gel electrophoresis was performed as previously described [4, 50]. Briefly, 4 µg of amoeba extract was separated on a 12 % SDS-polyacrylamide gel co-polymerized with 0.1 % gelatin. This was followed by incubation with 2.5 % Triton X-100 for 1 h, followed by incubation with 100 mM sodium acetate (pH 4.5), 1 % Triton X-100, and 20 mM DTT for 3 h at 37°C. Gels were stained with Coomassie to visualize cysteine peptidase activity.

### Determination of hemolytic activity

The hemolytic activity assay was performed as described by Biller et al. [3]. For the assay, trophozoites and erythrocytes were mixed at a ratio of 1:2000 (1.25 x 10^5^ amoebae with 2.5 x 10^8^ erythrocytes per ml NaPBS) and then incubated at 37 °C for 1 h. After incubation, the cells were sedimented, and the hemoglobin released in the supernatant was measured at 530 nm in a spectrophotometer. Separately incubated erythrocytes and trophozoites were used as negative controls. To determine 100 % hemoglobin release, 2.5×10^8^ erythrocytes were lysed in 1 ml of water. Experiments were performed at least three times in quadruplicate. Significance was determined by the Mann-Whitney *U* test.

### Western blot

To prepare amoeba extracts for Western blot, trophozoites were washed twice with NaPBS and sedimented by centrifugation at 400 x *g* for 2 min at 4 °C. To minimize proteolysis, 20 µM *trans*- Epoxysuccinyl-L-leucylamido(4-guanidino)butan (E64, Sigma-Aldrich, Merck, Taufkirchen, Germany) was added. For lysis the trophozoites were alternately flash frozen in liquid nitrogen for 4 times. The lysates were centrifuged at 40,000 x *g* for 1 h at 4°C. The supernatants contained PBS-soluble proteins. Pellets were washed twice in ice-cold NaPBS and solubilized in NaPBS supplemented with 1% Triton X-100. Extracts (50 µg/lane) were separated on 13% SDS-PAGE gels under reducing conditions. Proteins were transferred to nitrocellulose membranes by the wet blotting technique, with 25 mM Tris-HCl, 192 mM glycine, 1.3 mM SDS, pH 8.3, and 20 % methanol as the transfer buffer. For Western blot analysis, a primary anti-c-myc primary antibody (Sigma-Aldrich) at 1:1000 dilution and a secondary anti-mouse horseradish peroxidase (HRP) antibody (Sigma-Aldrich) at 1:5000 dilution was used. Blots were developed using GE Healthcare Amersham™ ECL Prime Western Blotting Detection Reagent (fisher scientific, Thermo Fisher Scientific, Schwerte, Germany).

### Immunofluorescence analyses

To localize the proteins to be analyzed, transfectants expressing the corresponding c-myc fusion proteins were harvested, washed 1 x with NaPBS, and resuspended in 1 ml NaPBS. After centrifugation at 400 x *g* at 4°C for 5 min, the supernatant was discarded, the cell sediment was resuspended in 1 ml of 4 % paraformaldehyde (PFA; Hatfield, PA, USA) in NaPBS, and the trophozoites were incubated for 30 min at RT on a rolling incubator in the NaPBS/4 % PFA solution. This and all subsequent incubation steps were performed on a rolling incubator. After centrifugation at 400 x *g* for 3 min, the supernatant was discarded, and the trophozoites were resuspended in 500 μl NaPBS/0.05 % saponin (or permeabilization of cell membranes (Sigma-Aldrich, Merck, Taufkirchen, Germany) and incubated for 5 minutes. After centrifugation at 400 x g for 3 minutes, the supernatant was discarded and the trophozoites were resuspended in 500 μl 50 μM ammonium chloride solution to block free aldehyde groups and incubated for an additional 15 min at RT. This was followed by two washing steps with 500 μl NaPBS/0.05 % saponin or 500 μl NaPBS (400 x g for 3 min) and an incubation for 10 min with 500 μl NaPBS/2 % FCS resuspended. An additional washing step was followed by a 1 h incubation at RT with the primary mouse anti-c-myc antibody (500 µl, 1:200, Sigma Aldrich). Further, three washing steps followed as described above. The cell sediments were then incubated in 500 μl NaPBS containing the secondary fluorescently labeled antibody (1:400, anti-mouse alexa fluor 488, Thermo Fisher Scientific, Bremen, Germany) for another 1 h rolling at RT.

For co-localization, an antibody targeting the cytoplasmic iron-containing superoxide dismutase (SOD; dilution 1:200; [51]) and anti-rabbit Alexa Fluor 594 antibody (dilution 1:400; Thermo Fisher Scientific, Bremen, Germany) were used. After washing the trophozoites again three times, the nuclei were stained by incubation with Hoechst-33342 (1:400; Invitrogen, Thermo Fisher Scientific, Bremen, Germany) diluted in 500 μl NaPBS for 10 min at RT. After a final washing step the trophozoites were resuspended in 50 μl NaPBS and could be stored in the dark at 4°C until analysis. Microscopy was performed using a *Zeiss Axio Imager M2* microscope and an *Olympus FluoView1000* confocal microscope.

### Supporting information

**S1 Table. RNAseq analyses of clone A1^np^ and A1^np^MP8-1^Si^ silencing transfectant.**

**S2 Table. RNAseq analyses of clone A1^np^ and A1^np^MP8-2^Si^ silencing transfectant.**

**S3 Table. RNAseq analyses of clone A1^np^ and A1^np^MP8-1+2^Si^ silencing transfectant.**

**S4 Table. RNAseq analyses of clone B2^p^ and B2^p^MP8-1^Si^ silencing transfectant.**

**S5 Table. RNAseq analyses of clone B2^p^ and B2^p^EhHP127^Si^ silencing transfectant.**

**S6 Table. RNAseq analyses of A1^np^pNC and A1^np^HP127^OE^ transfectants.**

**S7 Table. Comparison of the amino acid sequences of various leishmanolysins and invadolysins with EhMP8-1 and EhMP8-2. Identity in % is shown.**

**S8 Table. Oligonucleotides for qRT-PCR and for generation of pNC-GOI^Myc^.**

## Author Contributions

**Conceptualization**: Juliett Anders, Constantin König, Corinna Lender, Iris Bruchhaus

**Data curation:** Juliett Anders, Constantin König, Corinna Lender, Iris Bruchhaus

**Formal analysis:** Juliett Anders, Constantin König, Corinna Lender, Iris Bruchhaus

**Funding acquisition:** Constantin König, Iris Bruchhaus

**Investigation:** Juliett Anders, Constantin König, Corinna Lender, Iris Bruchhaus

**Methodology:** Juliett Anders, Constantin König, Corinna Lender, Arne Hellhund, Sarah Nehls, Ibrahim Shalabi, Barbara Honecker, Martin Meyer, Jenny Matthiesen, Dániel Cadar

**Resources:** Juliett Anders, Constantin König, Corinna Lender, Jenny Matthiesen, Martin Meyer

**Software:** Juliett Anders, Constantin König, Stephan Lorenzen, Nahla Galal Metwally, Iris Bruchhaus

**Supervision:** Iris Bruchhaus

**Visualization:** Arne Hellhund, Constantin König, Nahla Galal Metwally, Iris Bruchhaus

**Writing – original draft:** Iris Bruchhaus

**Writing – review & editing:** Juliett Anders, Constantin König, Corinna Lender, Barbara Honecker, Stephan Lorenzen, Jenny Matthiesen, Dániel Cadar, Thomas Roeder, Nahla Galal Metwally, Hannelore Lotter, Iris Bruchhaus

